# FOXO1 mitigates the SMAD3/FOXL2^C134W^ Transcriptomic Effect in a Model of Human Adult Granulosa Cell Tumor

**DOI:** 10.1101/2020.12.15.422901

**Authors:** Christian Secchi, Paola Benaglio, Francesca Mulas, Martina Belli, Dwayne Stupack, Shunichi Shimasaki

## Abstract

**Background:** Adult granulosa cell tumor (aGCT) is a rare type of stromal cell malignant cancer of the ovary characterized by elevated estrogen levels. aGCTs ubiquitously harbor a somatic mutation in *FOXL2* gene, Cys134Trp (c.402C<G); however, the general molecular effect of this mutation and its putative pathogenic role in aGCT tumorigenesis is not completely understood. We previously studied the role of FOXL2^C134W^, its partner SMAD3 and its antagonist FOXO1 in cellular models of aGCT.

**Methods:** In this work, seeking more comprehensive profiling of FOXL2^C134W^ transcriptomic effects, we performed an RNA-seq analysis comparing the effect of FOXL2^WT^/SMAD3 and FOXL2^C143W^/SMAD3 overexpression in an established human GC line (HGrC1), which is not luteinized, and bears normal alleles of FOXL2.

**Results:** Our data shows that FOXL2^C143W^/SMAD3 overexpression alters the expression of 717 genes. These genes include known and novel FOXL2 targets (TGFB2, SMARCA4, HSPG2, MKI67, NFKBIA) and are enriched for neoplastic pathways (Proteoglycans in Cancer, Chromatin remodeling, Apoptosis, Tissue Morphogenesis, Tyrosine Kinase Receptors). We additionally expressed the FOXL2 antagonistic Forkhead protein, FOXO1. Surprisingly, overexpression of FOXO1 mitigated 40% of the altered genome-wide effects specifically related to FOXL2^C134W^, suggesting it can be a new target for aGCT treatment.

**Conclusions:** our transcriptomic data provide novel insights into potential genes (FOXO1 regulated) that could be used as biomarkers of efficacy in aGCT patients.

## 1. Background

Granulosa cell tumors (GCTs) are a rare form of stromal cell malignant cancer accounting for one in twenty cases of ovarian cancer. GCTs can occur as either adult or juvenile subtypes [1]. Commonly the adult form of GCTs is diagnosed at an early stage and can be treated surgically. However, these tumors often recur, and are predominantly fatal in case of recurrence [2–4]. Current research is focused on discovering new molecular markers to predict the progression of aGCTs.

Established cancer markers have no prognostic significance for this type of tumor. Indeed, while many types of tumors show prognostic induction of oncogenes or inactivation of tumor suppressor genes, aGCTs reveal a modestly impacted genome. The immunohistochemical evaluation of different oncogenes and tumor suppressors (*e.g.,* MYC, CDKN1A, ERBB2 and TP53) failed to detect a correlation with patient outcome [5, 6]. Early studies suggested that FOXL2^C134W^ is potentially involved in the onset of aGCTs [7–12], and it is still matter of investigation. More than 400 tumors from aGCT patients of diverse ethnicities [13] showed only one single, specific mutation in the Forkhead box L2 transcription factor gene FOXL2: Cys134Trp (c.402C<G) [14]. The mutation is not present in other cancers. FOXL2 is expressed in the ovarian granulosa cells (GCs) and plays a key role during the development of female reproduction system and in its maintenance [15–17]. Potential FOXL2 targets revealed by ChIP and RT-PCR implicated its involvement in follicular and craniofacial development [18–20]. It has been also shown that FOXL2 affects collagen synthesis [21, 22]. Nicol *et al.* carried out the first ChIP-seq on mouse FOXL2^WT^ in order to find its direct target [23]. In 2011, Benayoun *et al*. demonstrated that FOXL2 plays a key role in the homeostasis of GCs and its failure leads to ovarian aging and tumorigenesis [24]. This mutation hyperactivates aromatase and downregulates GC apoptosis pathways [8]. Whole transcriptome analysis of 10 aGCT patient samples showed that the FOXL2^C134W^ acts as a hypomorphic mutant on the normally FOXL2^WT^ activated genes [25]. Another transcriptomic study on several stage 1 and stage 3 aGCT samples identified 24 genes whose expression significantly fluctuated between stages [26]. More recently, whole genome sequencing of a large aGCTs cohort indicated *TP53* and *DICER1* as potential drivers in these tumors and found a higher mutational burden in recurrent tumors, as compared to primary AGCTs [27]. However, these molecular events are not frequent as FOXL2^C134W^. Very few publications investigated the FOXL2^C134W^ pathogenicity from a transcriptional perspective. Notably, Carles *et. al.* developed an inducible FOXL2^C134W^ stable luteinized cell line (SVOG3e) [28] which demonstrated that FOXL2^C134W^ mutation may precisely alter DNA binding specificity. The study revealed low numbers of target genes and the authors noted the SVOG3e cell line may not respond appropriately to FOXL2^C134W^. This may be possibly due to the absence of the needed partner SMAD3. Indeed, it is well known that SMAD3, a central element of the TGFβ pathway, is essential for the FOXL2^C143W^ activity [9, 29–36]. In line with this, a recent study by Weis-Banke S.E. *et al*. [37], showed that FOXL2^C134W^ and the TGFβ pathway are together two important potential therapeutic targets. Indeed, using FOXL2^C134W^ overexpression and silencing molecular approaches in a non-luteinized GC cell line (HGrC1) treated with TGFβ, they showed that SMAD4 is also an important FOXL2^C134W^ partner and the TGFβ/FOXL2^C134W^ molecular events trigger the expression of oncogenic, EMT, and stemness pathways in an aGCT model [37]. It is well accepted that TGFβ has an important role in tumor progression and involves SMAD mediators that control expression of hundreds of genes in different ways in diverse contexts and physiologies [38, 39] although additional investigations are needed to find its specific action(s) in aGCTs. To further demonstrate the role of SMAD3 as a molecular partner of FOXL2^C143W^ in aGCT progression on a genome-wide level, we carried out RNA-seq analysis in HGrC1 overexpressing FOXL2^C143W^ together with SMAD3, compared to FOXL2^WT^/SMAD3 as a control. We further explored the FOXL2 antagonistic tumor-suppressive transcription factor FOXO1 [35] to study its influence on the SMAD3/FOXL2^C134W^ induced alterations. FOXO factors are considered as tumor suppressors for their parts in the initiation of apoptotic process and in the cell cycle arrest [40]. We report that FOXL2^C134W^ over-expression was associated with altered signaling pathways previously recognized by transcriptomic studies from GC *in situ* [11, 37], and additionally identified novel candidate FOXL2^C134W^ targets. Furthermore, we observed that FOXO1 strongly mitigated the overall FOXL2^C134W^ action with important implications for both aGCT onset and future therapeutic development.

## 2. Materials and Methods

### 2.1. Plasmids and reagents

Expression plasmid encoding N-terminally Flag-tagged hFOXO1-3A (#13508) was purchased from Addgene (Cambridge, MA, pCDNA3 backbone). FOXO1-3A has Ala residue substitutions at Thr-24, Ser-256 and Ser-319 and is thus constitutively active. The usage of this mutant was based on the fact that protein remains in the nucleus due to the inability of insulin/growth factor signaling to phosphorylate the mutated residues. Like FOXL2 wild type (FOXL2^WT^), the FOXL2^C134W^ mutant is also localized in the nucleus [7] and thus FOXO1-3A is the best construct for our transcriptomic study. N-terminally Flag-tagged human FOXL2wt (pCS2+ backbone) was kindly given by Dr. Louise Bilezikjian [33] and its mutant (FOXL2^C134W^) was made in our laboratory as previously reported [41]. The expression plasmids encoding human SMAD3 were kindly provided by Drs. Kohei Miyazono [42] and Louise Bilezikjian [33].

### 2.2. HGrC1 cell culture

HGrC1 cells were kindly provided by Drs. T. Nakamura and A. Iwase, and their properties and derivation described earlier [43]. Notably, these cells bear two normal alleles of the FOXL2 gene [34]. Cells were cultured in a standard incubator at 37°C in DMEM-F12 medium (catalog #12400-024, Thermo Fisher Scientific, Waltham, MA) complemented with antibiotics (penicillin and streptomycin) and 10% FBS (#F6178, Sigma-Aldrich). HGrC1 cells at passage 14 were used for all the experiments in this study. HGrC1 cells seeded into 6-well tissue culture plates (1 x 10^6^ cells/well). Medium was replaced after 24 hours with serum-free DMEM-F12 (complemented with antibiotics), and transfection was performed using the above described plasmids for 4 hours with Lipofectamine 3000 reagents (catalog #L3000008, Thermo Fisher Scientific) following manufacturer’s procedures. After 4h, the medium was replaced with new serum free DMEM-F12 and transfected cells were cultured at 37°C for 24 hours.

### 2.3. RNA-seq data generation

After 4 hours transfection and additional culture for 24 hours in serum free DMEM-F12, HGrC1 cells (15 samples including 3 replicates for 5 conditions) were lysed using TRIzol reagent (catalog #15596026, Thermo Fisher Scientific) and RNA was extracted with Direct-zol RNA MiniPrep kit (#R2052, Zymo Research, Irvine, CA). RNA Quality control was performed by Agilent TapeStation. RNA-seq libraries were prepared using the TrueSeq Stranded mRNA Library Prep (Illumina, catalog #20020595) and sequenced using an illumina HiSeq4000 Platform using 75-bp single-end reads, to an average of 22.9 +/− 3.4 million reads per sample.

### 2.4. RNA-seq bioinformatics analyses

Differential expression analysis was performed with RNA-seq [44]. To quantify transcript abundance for each sample we used salmon (1.1.0) [45] with default parameters and using hg38 set as the reference genome for reads mapping. The data was subsequently imported in R using the tximport package [46] and processed with the DESeq2 package [47]. A total of 37,787 genes were analyzed. For Principal Component Analysis we normalized the raw transcript counts using *variance stabilized transformation* (*vst* function from DESeq2) and applied the *plotPCA* function from DESeq2, using the default 500 most variable genes to compute the principal components. All heatmaps were generated using vst expression values and the heatmap package in R, excluding genes from over-expressed vectors. Differential expression analysis was performed using DESeq2 comparing different pairs of conditions, and a threshold of Benjamini-Hochberg corrected p-value of 0.05 was chosen to define differentially expressed genes for each comparison. To estimate the fraction of genes with FOXL2 binding sites we used two publicly available human FOXL2 ChIP-seq dataset: one from TGFβ-treated HGrC1 cells ectopically expressing FOXL2 or FOXL2^C134W^ [37] and the other from an immortalized granulosa cell line (SVOG3e) with inducible expression of V5-tagged FOXL2^C134W^ protein [28]. For the first one (ChIP1), BAM files from FOXL2^C134W^ and the negative empty vector control were obtained from the authors and peaks were called using MACS2 [48] callpeak with default setting and using q-value cutoff of peak detection of 0.01 (-q 0.01). For the second (ChIP2), we obtained the peak coordinates from GEO (GSE126171) from 4 replicates at 12hrs induction of V5-tagged FOXL2^C134W^ and merged peaks coordinates of the replicates. Peaks BED files from the above ChIP-Seq were intersected with a +/− 5kb window from the transcription start site (TSS) of human genes (genecode v35 and genecode v19 respectively for ChIP1 and ChIP2) to identify genes with FOXL2^C134W^ binding around their promoter region (n= 9,939 and n=11,697, respectively). Enrichment for genes with FOXL2 binding was calculated using Fisher’s exact test in R, comparing DEGs versus non-DEGs.

### 2.5. Gene Ontology Enrichment analyses

For Gene Ontology Enrichment (GO) analyses, we used Metascape [49] with default parameters. To reduce the confounding effect on gene expression derived from variable overexpression of the proteins from the plasmids across samples, we removed the terms that were associated with the expression of FOXL2 and/or SMAD3. In order to do so, we performed Gene Set Enrichment Analysis (GSEA) and identified gene ontology terms significantly associated with FOXL2 and/or SMAD3. In detail, the GSEA tool (http://www.broad.mit.edu/gsea) for continuous phenotypes was used to identify gene annotation categories correlated with the genes of interest. For each gene of interest (FOXL2 or SMAD3), GSEA was run with the expression of that gene across samples set as the “continuous phenotype”; significant gene annotation sets with activity coordinated to the select gene were identified with adjusted P-value < 0.25. These annotation terms were removed from the results obtained with Metascape, *i.e.* from each list of terms enriched for differentially expressed genes throughout this investigation.

### 2.7. Data Accession

Datasets generated in this study will be made available in the NCBI GEO repository at the time of publication.

## 3. Results

### 3.1. Transcriptomics analysis of the human granulosa HGrC1 cell in presence of FOXL2^C134W^

To investigate the transcriptional changes associated with the SMAD3/FOXL2^C134W^ program, and its interaction with FOXO1, HGrC1 cells transiently expressing SMAD3 and FOXL2^WT^/FOXL2^C134W^, with or without FOXO1-3A (FOXO1 hereafter), were profiled using RNA sequencing (Fig. 1A). It is notable that endogenous FOXL2 and FOXO1 are also detectable in these cells [34, 35, 43]. An initial Principal Component Analysis (PCA) was calculated on variance stabilized transformation (*vst*)-normalized expression values from the top 500 most variable genes across samples. The variation between samples was largely explained by the de novo gene expression, with samples of the same conditions clustering closer together, indicating good reproducibility between replicates (Fig. 1B). To identify genes selectively induced by FOXL2 or FOXL2^C134W^, we next assessed differential expression analysis between FOXL2^WT^ and controls, FOXL2^C134W^ and controls, and FOXL2^C134W^ and FOXL2^WT^, and identified a total of 452, 939 and 717 differential expressed genes (DEGs, q<0.05) genes respectively, many of which were shared among the different conditions (Fig. 1C). Of note, FOXL2^C134W^ induced more changes with respect to the overexpression of the wild type protein, possibly indicating an enhanced transcriptional capacity.

**Fig. 1.**
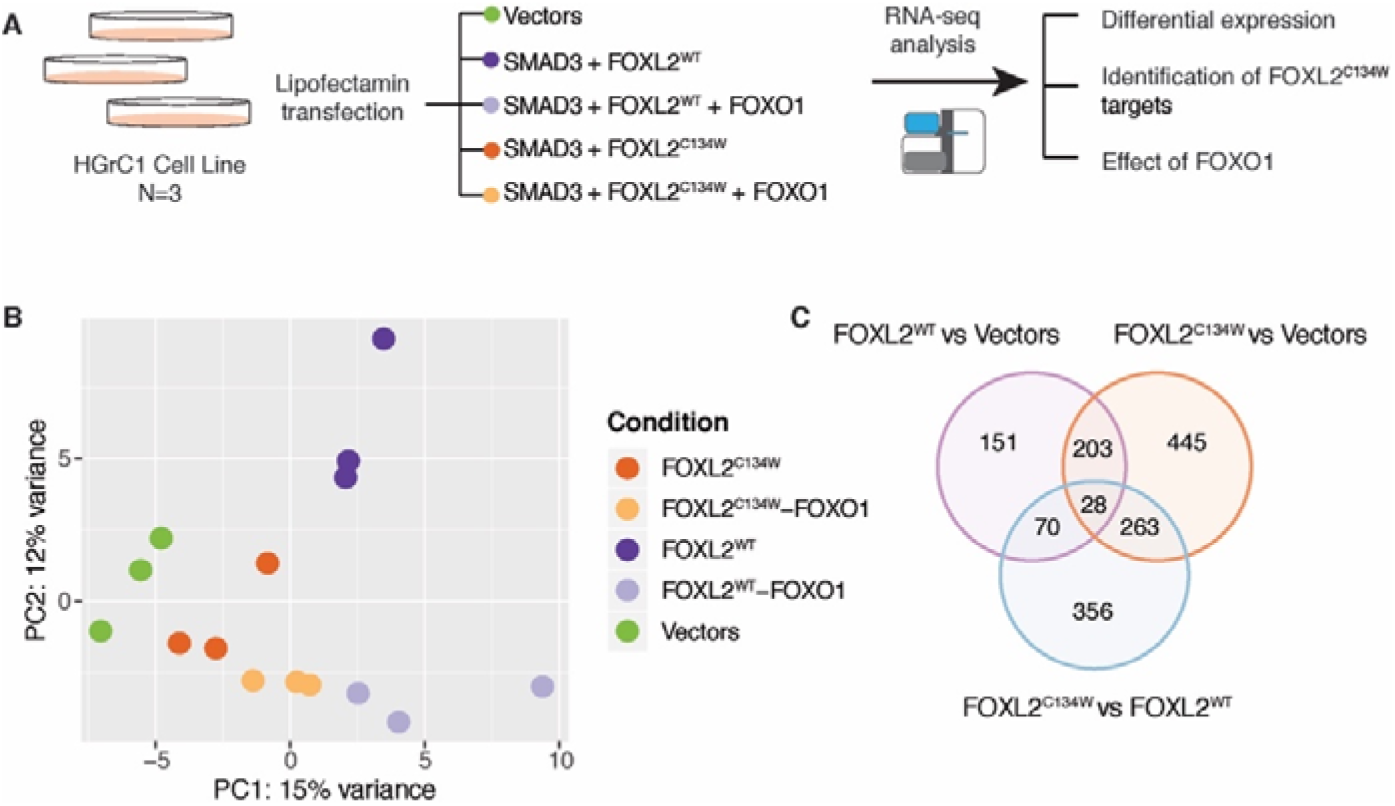
Transcriptome analysis of the human granulosa HGrC1 cell line under different transfection conditions. A) Schematic summary of study design. FOXL2^WT^/FOXL2^C134W^ were in pCS2+ expression plasmid constructs; FOXO1^3A^ in pCDNA3 expression construct to minimize promoter competition. B) Principal component 1 and 2 from all samples of the study, color-coded as indicated in the legend. Principal components were calculated on vst-normalized expression values from the top 500 most variable genes across samples. (SMAD3 was co-transfected with the other protein constructs per Methods but is omitted in the legend for brevity). C) Venn diagram showing the number of differential genes (DESeq, q<0.05) identified in 3 pairs of conditions: FOXL2^WT^ vs Vectors, FOXL2^C134W^ vs Vectors and FOXL2^C134W^ vs FOXL2^WT^.

### 3.2. Gene Ontology Analysis of DEGs of FOXL2^C134W^ vs FOXL2^WT^

To evaluate the functional significance of the genome-wide alterations in gene expression triggered by FOXL2^C134W^, we performed Gene Ontology (GO) analysis on DEGs, specifically focusing on the differences between FOXL2^C134W^ and FOXL2^WT^. The volcano plot (Fig. 2A) and heatmap analysis revealed significantly upregulated (*e.g.* SMARCA4, RRBP1, FLNA) and downregulated genes (e.g. NFKBIA, TAP1, CXCL8). Among the upregulated and downregulated differentially expressed genes (DEGs), many are associated with the (ECM) extracellular matrix (HSPG2, COL4A2, COL5A1) or cellular interaction with it (FLNA, FLNC), chromatin remodeling (SMARCA4, EP400, BAZ2A), and PI3K/AKT and inflammation pathways (AMIGO2, XBP1).

**Fig. 2.**
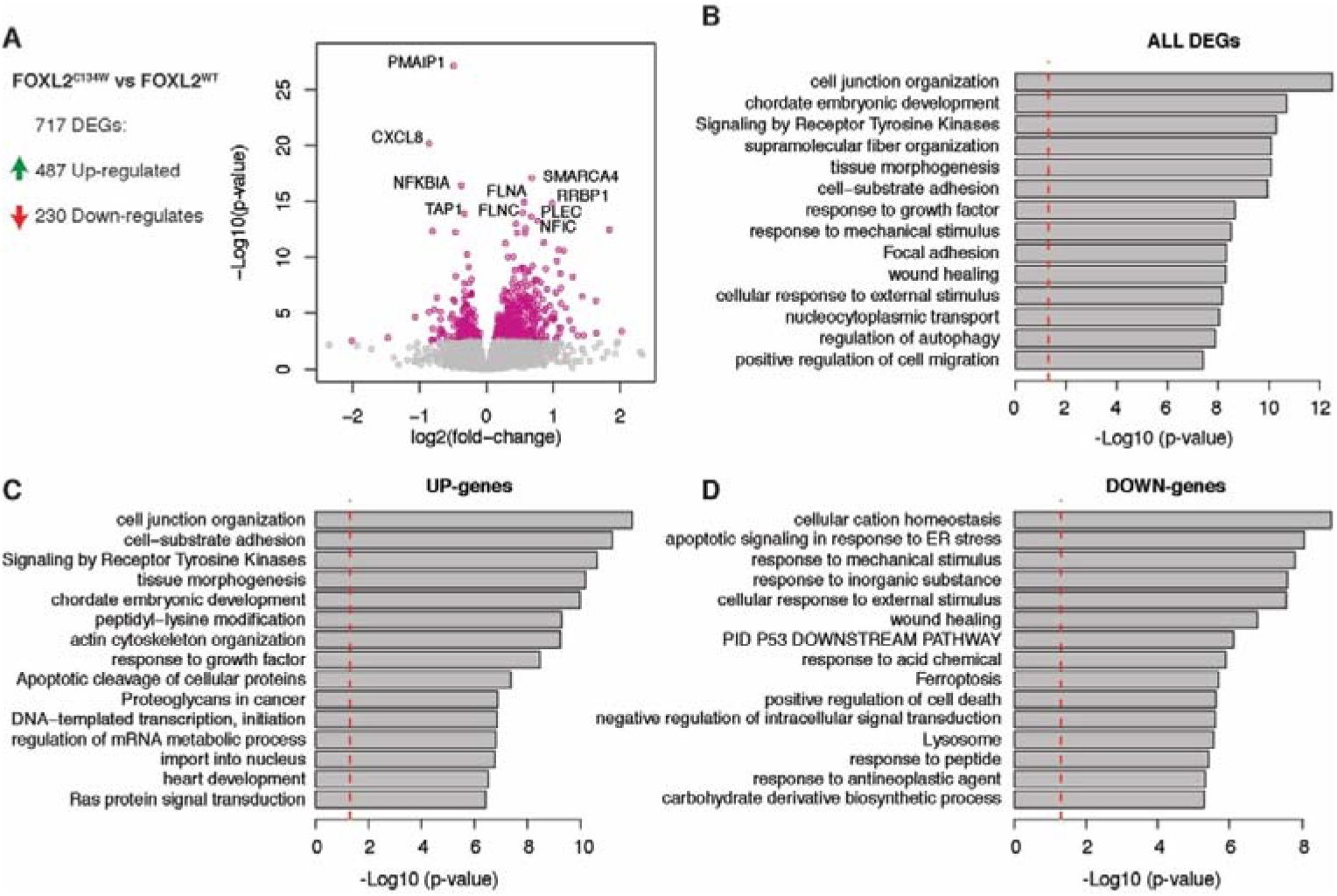
Gene Ontology enrichment analysis of differentially expressed genes (DEGs) between FOXL2^C134W^ and FOXL2^WT^. A) Volcano plot showing in pink DEGs between FOXL2^C134W^ and FOXL2^WT^ (DESeq, q<0.05). The genes FOXL2 and SMAD3 were omitted from the plot. B-D) Gene ontology enrichment analysis using all DEGs (B, n=717), up-regulated genes in FOXL2^C134W^ transfected cells (C, n=487), and down-regulated genes in FOXL2^C134W^ transfected cells (D, n = 230). The red line indicates p-value cutoff for p<0.05. Gene ontology enrichment p-values were calculated using Metascape [49]. GO terms associated with expression of FOXL2 and/or SMAD3 were removed from the results.

Enrichment analysis [49] of pathways was performed on the 717 DEGs overall (Fig. 2B) as well as on 487 upregulated (Fig. 2C) and 230 downregulated genes (Fig. 2D). The overall and the upregulated genes were highly enriched for gene annotation categories from oncogenic pathways, including Signaling by Receptor Tyrosine Kinases, and Proteoglycans in Cancer, Tissue Morphogenesis and Proliferation (wound healing and positive regulation of cell migration). Interestingly, GO categories enriched for downregulated genes (Fig. 2D) included pathways often suppressed by oncogenes. These contained Wound Healing and the PID P53 Downstream pathways. Overall these data support the pathognomic role of FOXL2^C134W^/SMAD3 complex as a driver of specific and selective oncogenic molecular events.

### 3.3. Identification of direct FOXL2 targets among DEGs between FOXL2^C134W^ and FOXL2^WT^

To determine whether DEGs were direct targets of FOXL2^C134W^, we next annotated gene promoters using available FOXL2^C134W^ ChIP-seq from the recent studies by Weis-Banke S.E. *et al*. [37] (ChIP1) in TGFβ-treated HGrC1, and by Carles *et al.* [28], which included inducible expression of FOXL2^C134W^ in human SVOG3e cell line (ChIP2). The majority (44%-40%) of DEGs between FOXL2^C134W^ and FOXL2^WT^ had a FOXL2 binding site at their promoters (313 out of 717 using the HGrC1 ChIP-seq and 291 using the SVOG3e ChIP-Seq), which was a significantly higher proportion than in non-DEGs (Fisher’s exact test, Fig. 3A overall p= 9.8×10^−51^; upregulated p=3.7×10^−37^; and downregulated p= 5.2×10^−15^; Fig. 3B overall p= 9.9×10^−39^; upregulated p=6.4×10^−26^; and downregulated p= 3.8×10^−14^). The presence of these binding sites suggests that the changes in gene expression in our system are due in large extent to differences in FOXL2 binding. To identify those targets differentially modified by FOXL2^C134W^, we focused on the subsets of upregulated and downregulated genes where the expression in FOXL2^C134W^ conditions was in the same direction of FOXL2^WT^ with respect to the vector. We selected the genes that were up or down in both mutant and wild-type conditions, reasoning that the approach is more robust at avoiding possible artifacts, and likely easier to interpret from a mechanistic point of view. There were 60 up-regulated genes and 82 down-regulated genes that satisfied these criteria (Fig. 3C, D). Among them were known targets of FOXL2, but also novel targets. Fig. 3C and 3D also show the comparisons between our outcomes and the two recent FOXL2 ChIP-Seq analysis [28, 37]. The comparative exploration indicated that our identified genes are direct FOXL2 targets [37] (ChIP1, 24 upregulated and 39 downregulated genes) and [28] (ChIP2, 22 upregulated and 47 downregulated genes). We also observed enrichment of genes associated with the tumorigenesis and metastasis (TGFB2, TGB1|1, EGR1, KLF4, NFKBIA, SDC4, ZYX) and GO categories that include oncogenic regulating pathways as *P53 signaling, NF-kappaB*, *Tyrosine Kinase Signaling*, and *Cytoskeleto*n. To compare our findings with those from overexpression of FOXL2^C134W^ in the human SVOG3e cell line [28], we intersected our list of 939 DEGs of FOXL2^C134W^ vs. Vector with the corresponding published list of 471 DEGs, and identified 90 common genes. Among these genes, we found TGB2, EGR1, NFKBIZ, SDC, and SDCBD which are homologues genes to the obtained ones in our previous analysis (Fig. 3C and 3D). Gene Ontology analysis of these 90 genes revealed cancer-related regulatory pathways.

**Fig. 3.**
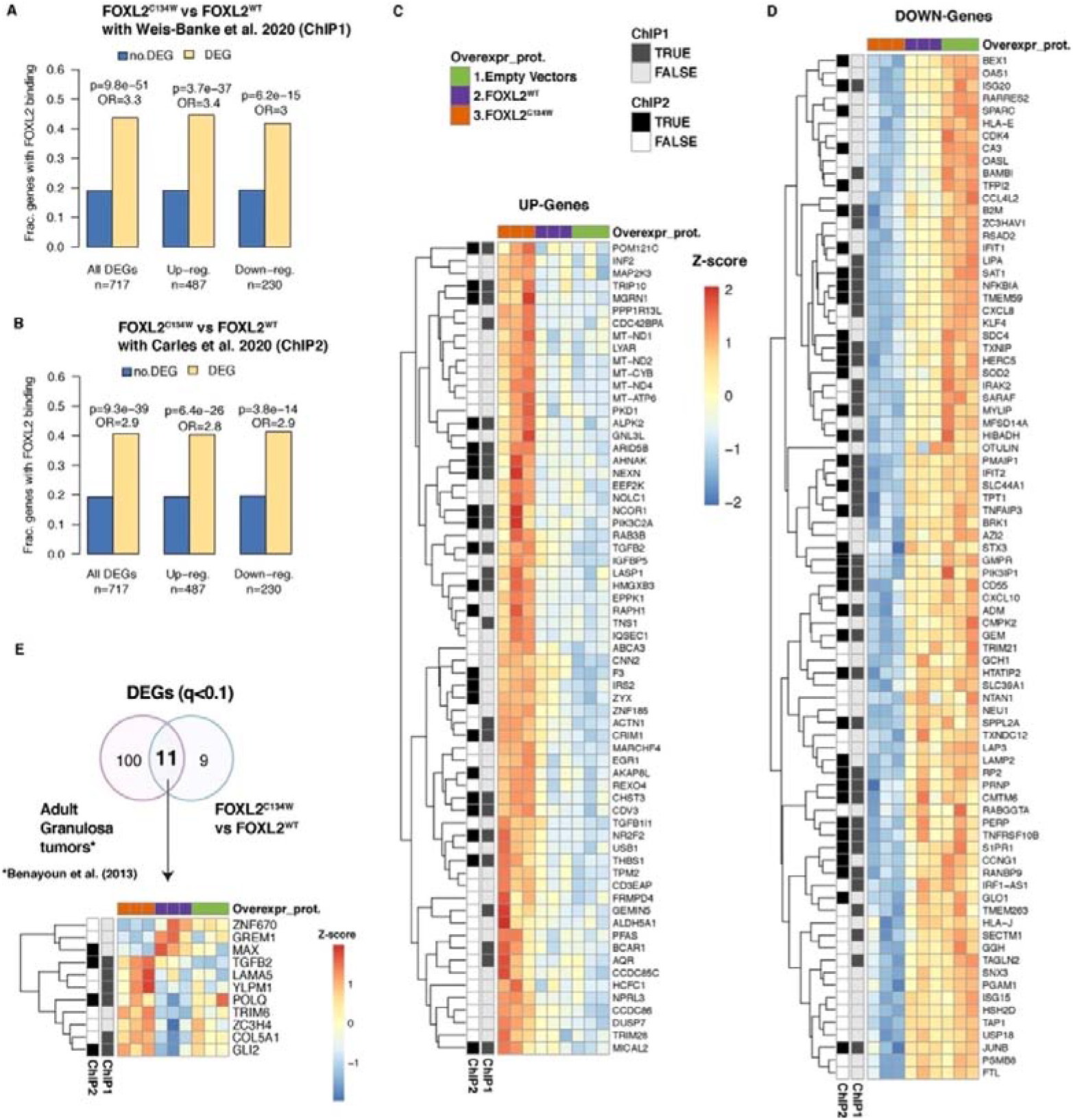
Identification of direct FOXL2 targets among DEGs between FOXL2^C134W^ and FOXL2^WT^. A) Enrichment of FOXL2 binding in DEGs between FOXL2^C134W^ and FOXL2^WT^ with respect to non-DEGs using FOXL2^C134W^ ChIP-seq binding site in TGFβ-treated HGrC1 (ChIP1) [37] and another B) using FOXL2^C134W^ ChIP-seq binding site in SVOG3e cell line (ChIP 2) [28]. P-values and odds ratios (OR) are from two-tailed Fisher’s exact test. C) Heatmap of the 60 up-regulated genes and D) Heatmap of the 82 down-regulated genes between FOXL2^C134W^ and FOXL2^WT^. Genes are ordered by p-value (DESeq) and by the satisfaction of homogeneity criteria where the expression in FOXL2^C134W^ conditions was in the same direction of FOXL2^WT^ with respect to the vector. For each gene, expression values were z-score normalized across samples. Genes are annotated for presence or absence of a FOXL2^C134W^ ChIP-seq binding site in TGFβ-treated HGrC1 (ChIP1) [37] and a FOXL2^C134W^ ChIP-seq binding site in a SVOG3e cell line (ChIP 2) [28]. E) Venn diagram (upper panel) showing the number of differential genes (q<0.1) identified in FOXL2^C134W^ vs FOXL2^WT^ (20 DEGs) from the present study and from adult granulosa cell tumor patient samples vs control samples (111 DEGs) previously published [25]. The lower panel shows heatmap of the common identified 11 genes. For each gene, expression values were z-score normalized across samples. Genes are annotated for presence or absence of a FOXL2^C134W^ ChIP-seq binding site in TGFβ-treated HGrC1 (ChIP1) [37] and a FOXL2^C134W^ ChIP-seq binding site in a SVOG3e cell line (ChIP 2) [28].

To complete our comparative analysis with the most recent transcriptomic data, we also intersected our FOXL2^C134W^ and FOXL2^WT^ DEGs list (717 genes) to the those DEGs in Weis-Banke S.E. *et al*. In addition, 29 common genes identified via Chip-seq data [37] were also compared to the DEGs. Analysis of these genes showed an involvement reproduction and tumorigenesis-related GO pathways like cell-adhesion regulation and negative regulation of wound healing. Finally, comparing our FOXL2^C134W^ and FOXL2^WT^ DEGs (717 genes) to transcriptomic data from aGTC patients published by Benayon *et al*. [25], we were able to categorize 11 genes (Fig. 3E), of which at least 3 (TGFB2, POLQ, GLI2) are *bona fide* FOXL2 targets [28, 37] (Fig. 3E). TGFB2 is the most consistent FOXL2^C134W^-modulated candidate and was present as target and upregulated genes in every analysis performed. These global results of our transcriptomic support the putative key role of FOXL2^C134W^ in aGCT tumorigenesis playing as regulator of oncogene (TGFB2, GLI2, EGR1) and as modulator of ECM genes (HSPG2, SDC4, COL5A1) and chromatin/nucleic acid remodeling (SMARCA4, BAZ2A).

### 3.4. FOXO1 overexpression mitigates FOXL2^C134W^ gene expression profile

We next assessed the effect of FOXO1 overexpression in addition to FOXL2^C134W^ on gene expression profiles. Interestingly, the number of DEGs between FOXL2^C134W^ in the presence of FOXO1, and FOXL2^WT^ were much lower (230) relative to the number of DEGs between FOXL2^C134W^ and FOXL2^WT^ (717, of which 130 in common), suggesting that the transcriptional changes of FOXL2^C134W^ compared to FOXL2^WT^ are largely mitigated by the co-expression of FOXO1 (Fig. 4A). This notion is supported by comparing the overall effect size for DEG between FOXL2^C134W^ and FOXL2^WT^ (n=717), with or without FOXO1 (Fig. 4B). While the log2FC are generally consistent across the two comparisons (FOXL2^C134W^ + FOXO1 vs FOXL2^WT^ and FOXL2^C134W^ vs FOXL2^WT^), the linear regression slope ß was 0.58 (less than 1), suggesting that up- or down-regulation of genes relative to the FOXL2^WT^ is attenuated by approximately 40% in presence of FOXO1. To further identify potential targets of FOXL2^C134W^ that are significantly modulated by FOXO1, we identified DEGs between FOXL2^C134W^ + FOXO1 and FOXL2^C134W^ and obtained 58 genes, of which 39 were in common with those between FOXL2^C134W^ and FOXL2^WT^ (Fig. 4C). These 39 genes (Fig. 4D), were downregulated in the FOXL2^C134W^ + FOXO1 vs FOXL2^C134W^ comparison, and vice versa. These genes were also compared against FOXL2 ChIP-seq from the recent studies by Weis-Banke S.E. *et al*. [37] and by Carles *et al.* [28]. Among these, 10 members were *bona fide* targets (including HSPG2, COL4A2 and BAZ2A). In particular, we identified the GO peptidyl-lysine modification pathway, implying that FOXO1 acts upon the chromatin remodeling triggered by FOXL2^C134W^. Moreover, other genes recognized in the ECM regulation were detected (HSPG2, COL4A2, COL5A1, COL18A1, DLG5). These data build on our prior identification of FOXO1 as a modulator of FOXL2 and provide the first molecular basis for FOXO1 in aGCT reprogramming.

**Fig. 4.**
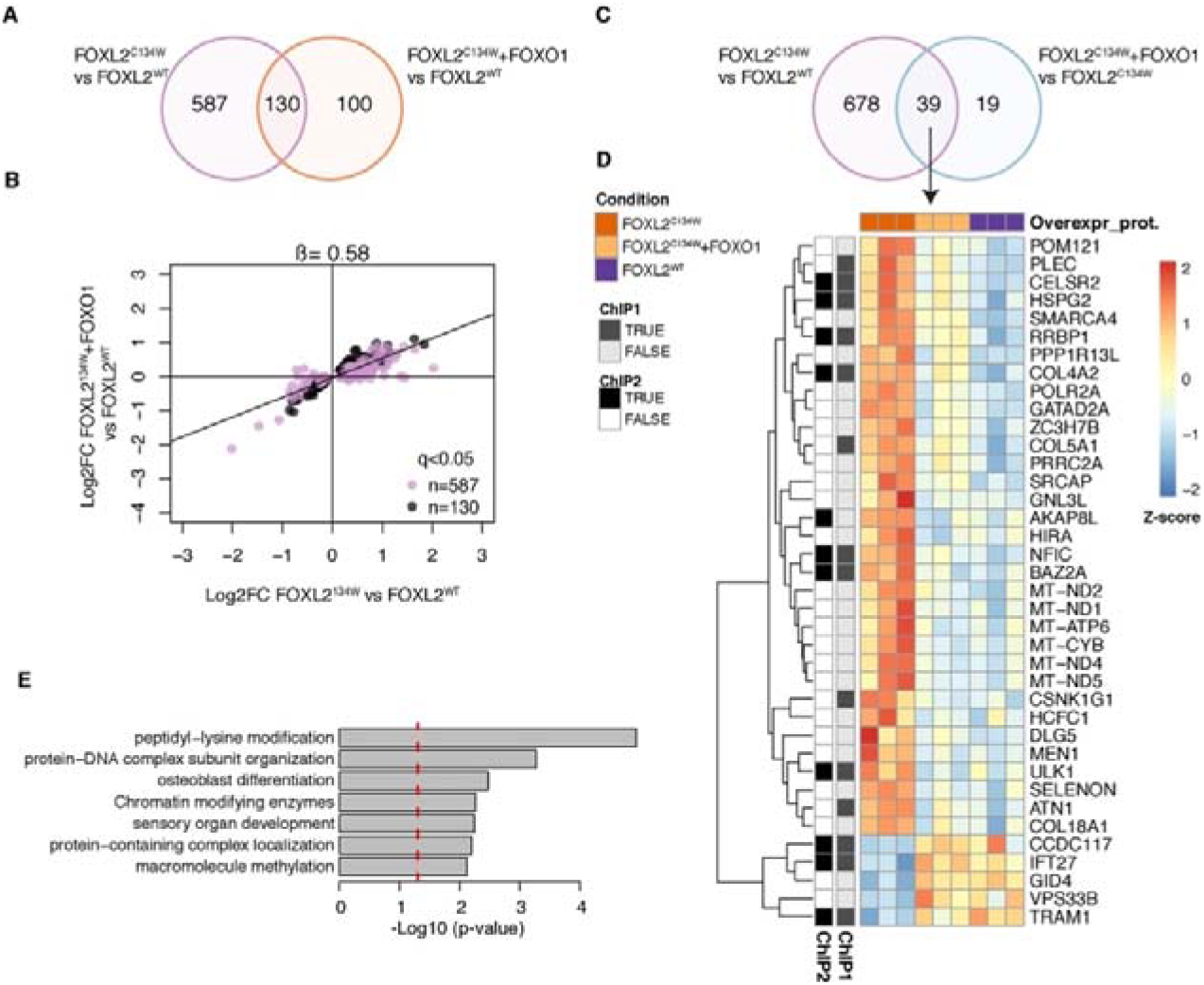
Effect of FOXO1 overexpression on FOXL2^C134W^ – modified genes. A) Venn diagram showing the number of differential genes (DESeq, q<0.05) identified in FOXL2^C134W^ vs FOXL2^WT^ and FOXL2^C134W^ + FOXO1 vs FOXL2^WT^. B) Scatter plot of the effect sizes (log2-fold change) of DEGs between FOXL2^C134W^ and FOXL2^WT^, with (y-axis) and without (x-axis) the addition of FOXO1. Genes that were differentially expressed also when adding FOXO1^3A^ are shown in black (n=130), while genes that were differentially expressed only without FOXO1^3A^ addition are shown in purple (n=587). The slope (ß) of the linear relationship between the two effect sizes is shown on top of the plot. C) Venn diagram showing the number of differential genes (DESeq, q<0.05) identified in FOXL2^C134W^ vs FOXL2^WT^ and FOXL2^C134W^ + FOXO1 vs FOXL2^C134W^. D) Heatmap with gene clustering of the 39 DEGs in both FOXL2^C134W^ vs FOXL2^WT^ and FOXL2^C134W^ + FOXO1 vs FOXL2^C134W^ comparisons. For each gene, expression values were z-score normalized across samples. Genes are annotated for presence or absence of a FOXL2^C134W^ ChIP-seq binding site in TGFβ-treated HGrC1 (ChIP1) [37] and a FOXL2^C134W^ ChIP-seq binding site in a SVOG3e cell line (ChIP 2) [28]. E) Metascape GO analysis of the 39 DEGs in both FOXL2^C134W^ vs FOXL2^WT^ and FOXL2^C134W^ + FOXO1 vs FOXL2^C134W^ comparisons. The red line indicates p-value cutoff for p<0.05. GO terms associated with expression of FOXL2 and/or SMAD3 were removed from the results (see Methods).

## 4. Discussion

In contrast to the rarer juvenile Granulosa Cell tumors, the bulk of granulosa cell tumors are the adult type (aGCT) which occur peri to post-menopausally [50–52]. Patients characteristically present elevated estrogen levels and postmenopausal bleeding [3, 53]. Almost all aGCTs bear the unique somatic mutation FOXL2^C134W^ [54], which appears essential to the aGCT development [7, 9, 14, 25, 55–58]. However, its pathogenic role in the aGCTs has been investigated only recently and the FOXL2^C134W^ transcriptional action in aGCTs is still currently studied. These investigations are not trivial, as transcriptomic studies on rare tumors - like aGCTs – face difficulties in the procurement good quality RNA samples. Control samples from healthy human granulosa cells without any previous treatment (HGC or hormones) are similarly limited. Accordingly, very few transcriptomic explorations based on aGCTs samples are present in the literature [25, 26, 28, 37]. Carles *et al.* developed an inducible FOXL2^C134W^ stable luteinized cell line (SVOG3e) [28] showing FOXL2^C134W^ can precisely modify DNA binding specificity. More recently Weis-Banke S.E. *et al*. [37] performed an interesting transcriptomic study essentially investigating FOXL2^C134W^ overexpression in presence of TGFβ on the same non-luteinized GC cell line (HGrC1) used in our study. They showed that mutant FOXL2 has an important partner not only in SMAD3, but also in SMAD4. This corroborates our previous hypothesis that the partnership between the FOXL2^C134W^ and TGFβ-pathway SMADs mediators is essential and specifically orchestrates the aGCT gene expression hallmarks [34, 35]. Here, we carried out a novel and complementary transcriptomic investigation overexpressing FOXL2^C143W^ in HGrC1 with its essential partner, SMAD3 [9, 29–36] to better understand its regulation and also avoiding other inevitable pleiotropic actions of the TGFβ stimulation (Fig.1). Gene Ontology (GO) analysis on the 717 DEGs (FOXL2^wt^ vs. FOXL2^C143W^) revealed a clear engagement of pathways and genes related to tumorigenesis: Tissue Morphogenesis and Proliferation (MKI67, MYH9, FLNA), Regulation of the Extracellular Matrix (ECM) and Proteoglycans in Cancer (HSPG2, FLNC, COL4A2, COL5A1), Chromatin Remodeling with the Peptidyl-Lysine modification (SMARCA4, EP400, BAZ2A), downregulation of apoptotic pathway (ATF3, ATF4, CDKN2D, CXCL8), and PI3K/AKT pathway (AMIGO2, XBP1) (Fig. 2).

Although MKI67, SMARCA4 and NF-kappaB pathway components have been described in aGCTs [5, 59–61] they have not previously been associated with FOXL2^C134W^ or implicated as potential direct targets. Importantly, a significant majority of the 717 DEGs had a FOXL2 binding site at their promoters: 313 were identified using the TGFβ treated-HGrC1 ChIP-seq [37] (annotated ChIP1) and 291 using the SVOG3e ChIP-Seq [28] (ChIP2). This suggests that the variations in gene expression in our system are related to difference in FOXL2 binding (Fig.3A and Fig 3B). We considered to be important the comparison with FOXL2 ChIPseq analysis from Weis-Banke S.E. *et al*. (ChIP1) since it was validated with the hybrid DNA– binding sites in RNA extracted from 4 aGCT patients and, therefore, can be easily referred to as the best *bona fide* enrichment for FOXL2 for the aGCT investigation in the current literature. A stringent selection of a subsets of 60 upregulated and 82 downregulated genes (Fig. 3C, Fig. 3D) in which the expression in FOXL2^C134W^ sample conditions was in the same direction of FOXL2^WT^ with respect to the vector allowed us to identify more specific targets: 17 up- and 27 down-genes annotated in both ChIP-seq analysis [28, 37]. Most of these targets (TGFB2, TGFB1|1, EGR1, KLF4, NFKBIA, SDC4, ZYX) are associated with neoplastic-events. Interestingly, 11 genes - whose 3 (TGFB2, GLI2, POLQ) are direct FOXL2 targets according previous ChIP-seq studies - were observed to be common between the FOXL2^C134W^ and FOXL2^WT^ DEGs (717 genes) and the transcriptomic data from aGTC patients published by Benayon *et al.* [25] (Fig. 3E). 90 genes have been identified in common between our study and that of Carles *et al.* [28] (FOXL2^C134W^ vs Vectors condition, not shown). 29 common genes were also identified in common with the Weis-Banke S.E. *et al*. [37] (FOXL2^C134W^ vs FOXL2^WT^, not shown). It was remarkable that the TGBF2 gene emerges as the strongest FOXL2^C134W^ modulated candidate since was present as target and upregulated genes in all the analysis carried out. This is consistent with the molecular relationship between FOXL2 and TGFβ family [13, 41, 62]. Indeed, FOXL2^C134W^ can trigger neoplastic events in granulosa cells by altering the TGFβ pathway [13]. Importantly, this finding implies that FOXL2 mutant may drive the steady induction of its key molecular SMAD partners by via a direct TGFB2 positive feedback loop. aGCT therapeutic approaches based on the inhibition of the TGFβ family member, Activin showed limited antitumoral activity. However, potentially clinically meaningful dose-related metabolic effects, including treatment of cancer cachexia, were observed that support further exploration of activin A inhibitors [63, 64]. Treatments based on the Activin inhibition and other novel TGFβ pathway targeting approaches – as strongly advocated by Weis-Banke S.E. *et al*.[37] and currently optimized for other indication [65–67] will be necessary to better determine their clinical meaning in aGCT [63, 64]. Interestingly, the chromatin remodeler SMARCA4 (BSGR1) - already recognized as an important player in cellular tumorigenesis [68, 69] and expressed in all aGCT samples [60] - has been directly associated with TGFβ pathway [70] supporting the assumption of FOXL2^C134W^ as main pathognomic orchestrator of aGCT. Our overall data are a novel contribution in deciphering how FOXL2^C134W^ coordinates aGCT tumorigenesis.

An important result in our study came from the transcriptomic assessment of the onco-suppressor FOXO1 effect in our system. FOXO1 is the most abundantly expressed member of FOXO gene family, and regulates vital cellular processes including the cell cycle, apoptosis, energy homeostasis and ROS catabolism [71]. Insulin signaling through the PI3K-AKT pathway is the main modulator of FOXO1 [72]. FOXO1 phosphorylation during PI3K-AKT signaling promotes its translocation from the nucleus to cytoplasm and its consequent inactivation and proteasomal degradation [73]. The activation of PI3K-AKT signaling is common across many tumor types [74, 75] and the inactivation of FOXO1 in response to PI3K-AKT represents a common mechanism by which neoplastic cells prevent apoptosis and physiological cell cycle arrest [76–78]. It is well known that AKT inhibits FOXO through direct phosphorylation [79, 80]. Notably, FOXO1 can be indirectly targeted *via* the action of AKT inhibitors [81–84]. Our data, depicted in Fig. 4A, revealed that ectopic expression of FOXO1 specifically and strongly moderated the overall effect of FOXL2^C134W^ compared to FOXL2^WT^, directly repressing the number of DEGs between FOXL2^C134W^ + FOXO1 and FOXL2^WT^. This parallels our previous findings where our data indicated the inhibitory action of FOXO1 to SMAD3/FOXL2^C134W^ activity [35]. We identified potential targets of FOXL2^C134W^ that are significantly modulated by FOXO1 (58 genes, of which 39 in common) in the DEGs between FOXL2^C134W^ + FOXO1 and FOXL2^C134W^ (Fig. 4C and 4D) confirming that FOXO1 attenuates DEGs genes that were upregulated/downregulated by FOXL2^C134W^. Finally, we identified 10 targets genes annotated in both ChIP-seq analysis [28, 37] (Fig. 4D). GO analysis of these genes (Fig. 4E) revealed FOXO1 modified similar pathways (peptidyl-lysine modification, chromatin remodeling, and ECM regulation) as those triggered by FOXL2^C134W^.

## 5. Conclusions

The comparative transcriptomic data presented in this investigation open new scenarios in understanding FOXL2^C134W^ mediate regulation of cell signaling, and illustrate: (i) that FOXL2^C134W^ can act to modulate important aGCT oncogenic pathways, including chromatin remodeling (SMARCA4, BAZ2A) and ECM regulation (HSPG2, COL4A2); (ii) the role of TGFB2, as upregulated effector and target gene in the aGCTs molecular occurrences; (iii) a potential molecular basis of emerging approaches targeting AKT/PI3K pathway [85–87]. Finally, our data provide novel insights into potential genes (FOXO1 regulated) that could be used as biomarkers of efficacy in these patients.

## Abbreviations

aGCT: adult-type granulosa cell tumor
DEGs: differentially expressed genes
DMEM-F12: Dulbecco’s modified Eagle medium – F12
FOXL2: forkhead box L2
FOXO1: forkhead box O1
GC: granulosa cell
GCT: granulosa cell tumor
mRNA: messenger RNA
PCR: polymerase chain reaction
RT-PCR: quantitative reverse transcription polymerase chain reaction
SBE: SMAD binding element
SEM: standard error of the mean

## Acknowledgment

The authors would like to thank Dr. Louise Bilezikjian for FOXL2^wt^ expression plasmid, Dr. Kohei Miyazono for providing us with the original hSMAD3 cDNA clones, Dr. Kristen Jepsen for the RNA-seq platform assistance and the generation of the library, Dr. Cloos for sharing BAM files of the ChIP-seq data from Weis-Banke *et al.*, 2020.

## Authors’ Contributions

Author Contributions: C.S. contributed to acquisition of the data. C.S., P.B., F.M. contributed to analysis of data. C.S. M.B. and S.S. contributed to study conception and design. C.S., P.B., F.M., D.S., and S.S. contributed to the interpretation of data and manuscript preparation.

